# Brain state and polarity dependent modulation of brain networks by transcranial direct current stimulation

**DOI:** 10.1101/179556

**Authors:** Lucia M. Li, Ines R. Violante, Rob Leech, Ewan Ross, Adam Hampshire, Alexander Opitz, John C. Rothwell, David W. Carmichael, David J. Sharp

## Abstract

Transcranial direct current stimulation (TDCS) has been widely used to improve cognitive function. However, current deficiencies in mechanistic understanding hinders wider applicability. To clarify its physiological effects, we acquired fMRI whilst simultaneously acquiring TDCS to the right inferior frontal gyrus (rIFG) of healthy human participants, a region involved in coordinating activity within brain networks. TDCS caused widespread modulation of network activity depending on brain state (‘rest’ or choice reaction time task) and polarity (anodal or cathodal). During task, TDCS increased salience network activation and default mode network deactivation, but had the opposite effect during ‘rest’. Furthermore, there was an interaction between brain state and TDCS polarity, with cathodal effects more pronounced during task performance and anodal effects more pronounced during ‘rest’. Overall, we show that rIFG TDCS produces brain state and polarity dependent effects within large-scale cognitive networks, in a manner that goes beyond predictions from the current literature.

## Introduction

Transcranial direct current stimulation (TDCS) has been extensively used in an attempt to modulate cognitive function in both healthy and disease populations (1). However, the behavioural results are variable and a recent meta-analysis concluded that single-session TDCS produces no effect on a range of cognitive tasks (2). This has fuelled scepticism about whether TDCS has any potential to modulate brain activity and cognitive function.

Interpreting the reasons for the varying behavioural effects of TDCS is limited by a lack of mechanistic understanding about its effects particularly at the level of the large scale brain networks. Cognitive function is mediated by the coordinated action of large scale brain networks (3,4) and therefore any cognitive effects of TDCS should have related alterations in brain network activity at this scale. The effects of TDCS can be influenced by the stimulation polarity and duration, and the relative effects on cognitive brain networks are largely unkown. It is therefore important to directly investigate the effects of TDCS on large-scale brain network activity, to help clarify the mechanism by which TDCS may influence behaviour.

Concurrent TDCS/functional MRI (fMRI) is an ideal method for studying the physiological effects of stimulation but, to date, such studies are few in number. Previous TDCS/fMRI studies have focused on the motor system. These have shown that that TDCS can modulate brain activity, as measured by haemodynamic changes, with effects seen remote from the site of stimulation measured by Blood Oxygenation Level Dependant (BOLD) functional MRI (5–9). Studies of the primary motor cortex (M1) suggest that anodal and cathodal TDCS have opposite effects on neuronal excitability (10). However, these motor studies cannot be extrapolated to explain effects on brain networks that support cognition. Moreover, the influence of polarity on cognitive network function and behaviour is poorly characterised, and there have been no studies focussing on how polarity interacts with underlying cognitive brain state.

Here we investigate the physiological effects of TDCS on large-scale brain networks that support cognition using simultaneous TDCS/fMRI. We employed a factorial design to manipulate both cognitive state (choice reaction task (CRT) or rest) and stimulation polarity (anodal or cathodal) (Figure 1). A protocol based on short stimulation durations (seconds) enabled a direct comparison between the effects of stimulation during different cognitive brain states, with different polarities and interactions between brain state and polarity.

**Figure 1:**
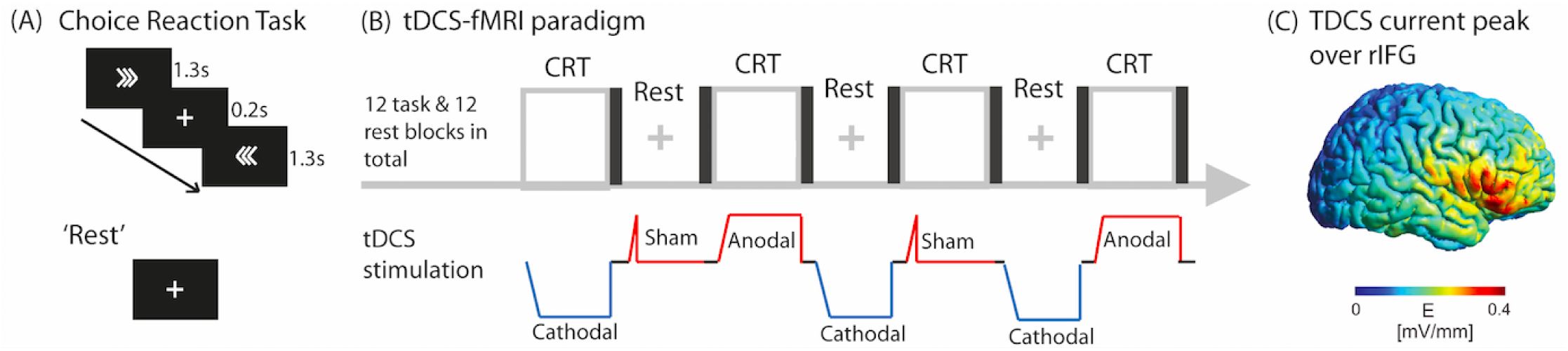
(A) Stimuli in the Choice Reaction Task. (B) The TDCS/fMRI paradigm, comprising 4 blocks each of CRT+anodal, CRT+cathodal, CRT+sham, ‘rest’+anodal, ‘rest’+cathodal and ‘rest’+sham TDCS. Each block was followed by a brief period of black screen and no stimulation. All participants underwent the paradigm 3 times. (C) Modelling showing peak current density over the rIFG.

We investigated the effects of stimulating the right inferior frontal gyrus (rIFG) (Figure 1C) because of its central role in cognitive function (11). This region and the underlying anterior insula form part of the salience/cingulo-opercular network (SN), which additionally comprises dorsal anterior cingulate cortex and pre-supplementary motor area (dACC/pre-SMA). Activity of the rIFG is seen in a range of cognitive contexts and the region is thought to influence activity in other cognitive regions, acting as a switch between different cognitive states (3,12,13), and influencing activity within the more extensive fronto-parietal control network (FPCN) as well as anti-correlated activity within the default mode network (DMN) (14). Hence, we tested the hypotheses that: 1) TDCS to the rIFG can modulate the function of intrinsic large-scale brain networks relevant to cognitive function; 2) the effects of TDCS will interact with cognitive brain state, and that 3) anodal and cathodal TDCS will have distinctive effects on cognitive network activity.

## Results

### The effects of transcranial direct current stimulation (TDCS) on brain activity are dependent on cognitive state

TDCS accentuated the patterns of activation and deactivation normally observed in each cognitive state. CRT performance compared to rest was characterized by increased BOLD response within the FPCN, including the SN, as well as primary sensory/motor cortices, bilateral thalami and basal ganglia, and decreased BOLD signal within the DMN (Figure 2A) (*SI Results*). Anodal and cathodal stimulation applied during CRT performance to the rIFG both increased activity within the dorsal anterior cingulate cortex /pre-supplementary motor area (dACC/pre-SMA) and lateral prefrontal regions. Cathodal TDCS was associated with an increased BOLD response within the SN, including the pars opercularis of the rIFG, as well as additional increases in bilateral frontal eye fields, bilateral basal ganglia, thalami and superior parietal activation (Figure 2Bi). Both anodal and cathodal TDCS decreased the BOLD response within medial parietal and occipital areas and medial pre-central gyrus (Figure 2Bii).

**Figure 2:**
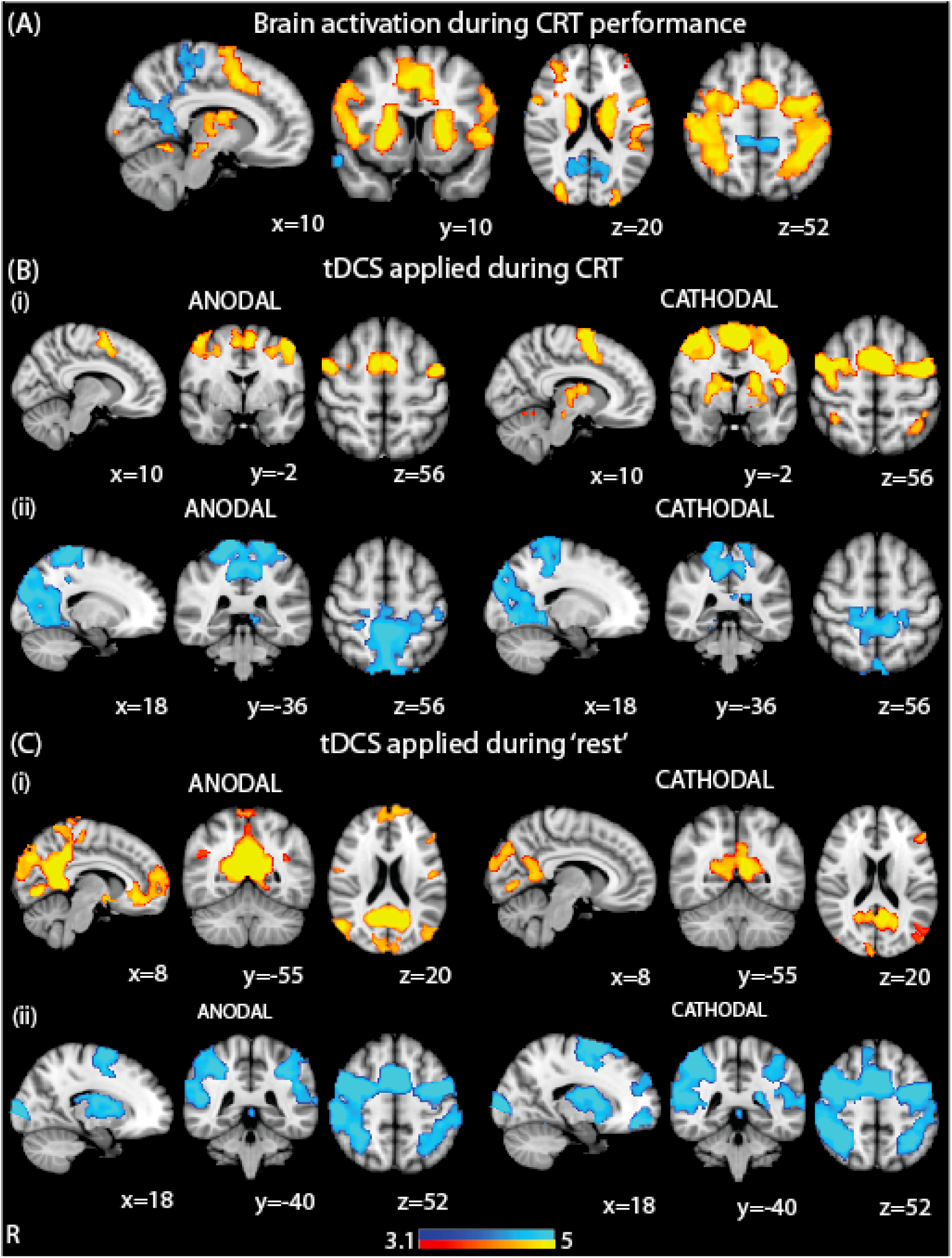
Accentuation of underlying patterns of brain activity with TDCS. (A) Overlay of brain activation and deactivation associated with CRT performance (with no TDCS). (B) Brain areas showing greater activation (i) and deactivation (ii) when TDCS is applied during CRT performance. (C) Brain areas showing greater activation (i) and deactivation (ii) when TDCS is applied during ‘rest’. Warm colours represent brain regions showing more activation and cool colours represent brain regions showing more deactivation. Results are superimposed on the MNI152 1mm brain template. Cluster corrected z=3.1, p<0.05.

In contrast, TDCS applied during ‘rest’ accentuated the activation of the DMN. Stimulation increased activity within the posterior cingulate cortex (PCC), the precuneus, inferior parietal regions and medial occipital areas. Anodal TDCS was associated with more extensive medial occipital and PCC activation as well as additional increases in the ventromedial prefrontal cortex (vmPFC) (Figure 2C1). Both types of stimulation showed a similar pattern of reduced activation within SN, FPCN and thalamic and basal ganglia regions (Figure 2Cii).

Both anodal and cathodal TDCS, during ‘rest’, produced increased BOLD signal in the rIFG close to the site of stimulation. However, widespread BOLD changes were observed mainly in brain areas anatomically remote from the cortical area being stimulate.

An ANOVA of the BOLD activity was performed with two factors, brain region (4 levels: dACC/preSMA, rIFG, PCC and vmPFC) and condition (4 levels: task+anodal, task+cathodal, ‘rest’+anodal, ‘rest’+cathodal). As well as an overall effect of condition (F(3,375)=5.20, p=0.0016) and brain region (F(3,374)=9.65, p<0.0001), there was an interaction between brain region and condition (F(9,375)=21.86, p<0.0001), which suggests that TDCS does not accentuate network activity uniformly across the network but has a more complex effect.

### Polarity dependent effects of transcranial direct current stimulation that interact with cognitive brain state

The effects of TDCS were also dependent on the polarity of stimulation. Directly contrasting cathodal and anodal TDCS showed polarity dependent effects of TDCS, which interacted with brain state (Figure 3). When applied during CRT performance, cathodal TDCS produced increased activity in the dACC/pre-SMA node of the SN, as well as left superior frontal and parietal areas, compared to anodal TDCS (Figure 3A). These areas overlapped with parts of the left FPCN, usually activated by CRT performance. Conversely, when applied during ‘rest’, anodal TDCS produced greater activation in the DMN compared to cathodal TDCS (Figure 3B). These polarity-dependent effects were only seen in brain regions where brain activity increased during TDCS i.e. the large decreases in brain activation produced by TDCS were not polarity dependent.

**Figure 3:**
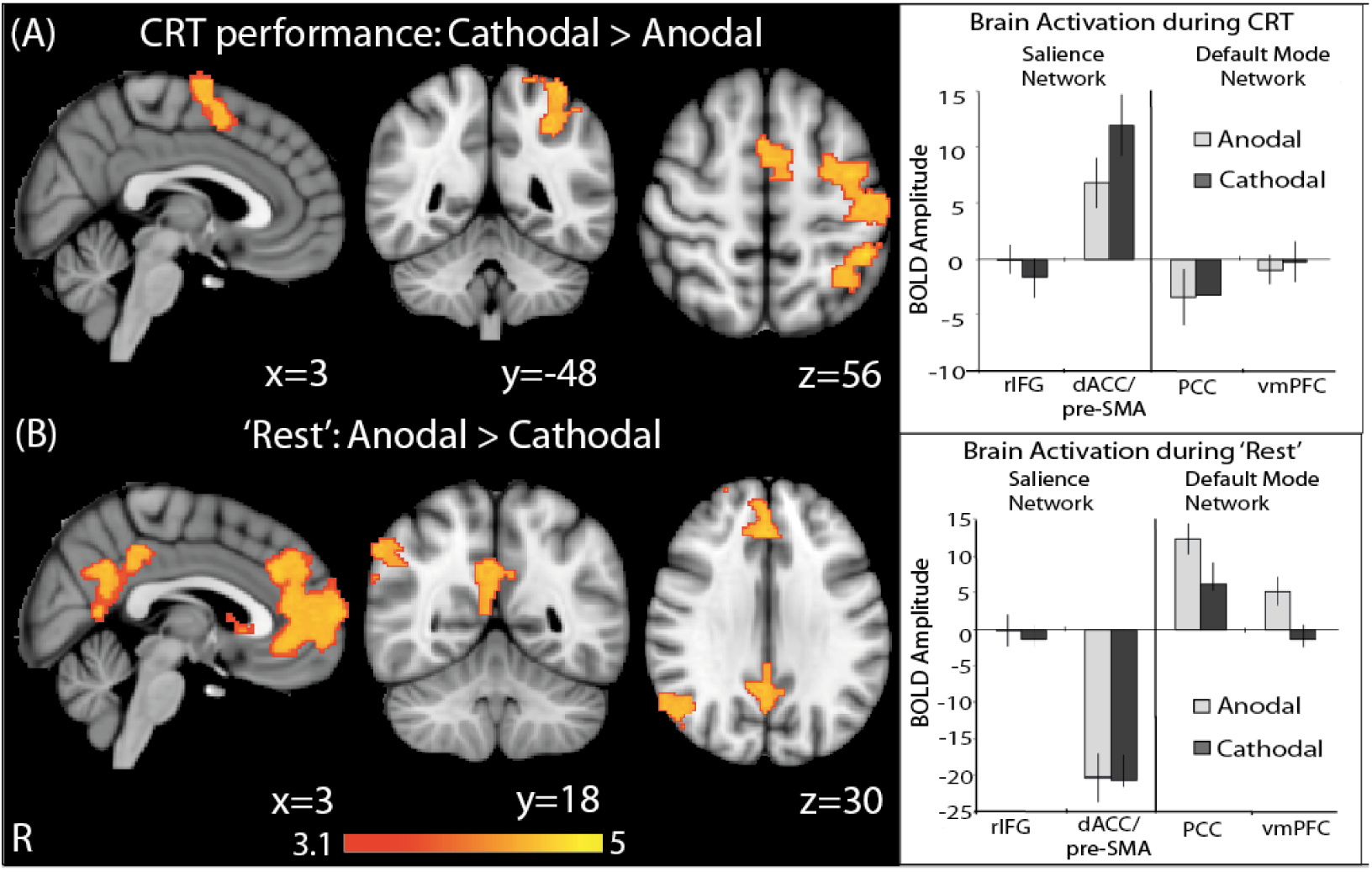
Differences in brain activation under anodal TDCS and cathodal TDCS. (A) Brain areas showing greater activation under cathodal than anodal TDCS during task. The accompanying bar chart shows mean activation in regions of interest within the Salience and Default Mode networks and are provided to demonstrate the change in BOLD magnitude with stimulation, as compared to task alone. (B) Brain areas showing greater activation under anodal than cathodal TDCS during ‘rest’. The accompanying bar chart shows mean activation in regions of interest within the Salience and Default Mode works and are provided to demonstrate the change in BOLD amplitude with stimulation, as compared to ‘rest’ without stimulation. Results are superimposed on the MNI152 1mm brain template. Cluster corrected z=3.1, p<0.05. Error bars denote S.E.M.

### Transcranial direct current stimulation modulates Salience Network connectivity

We next investigated whether TDCS modulated functional connectivity within intrinsic connectivity networks using psychophysiological interaction (PPI) analysis (15,16). Functional connectivity was investigated between nodes of the SN (the rIFG and dACC/pre-SMA) and DMN (the vmPFC and PCC). Cathodal TDCS applied during CRT performance produced increased functional connectivity between the dACC/pre-SMA (seed region) and the rIFG (Figure 4Ai).

**Figure 4:**
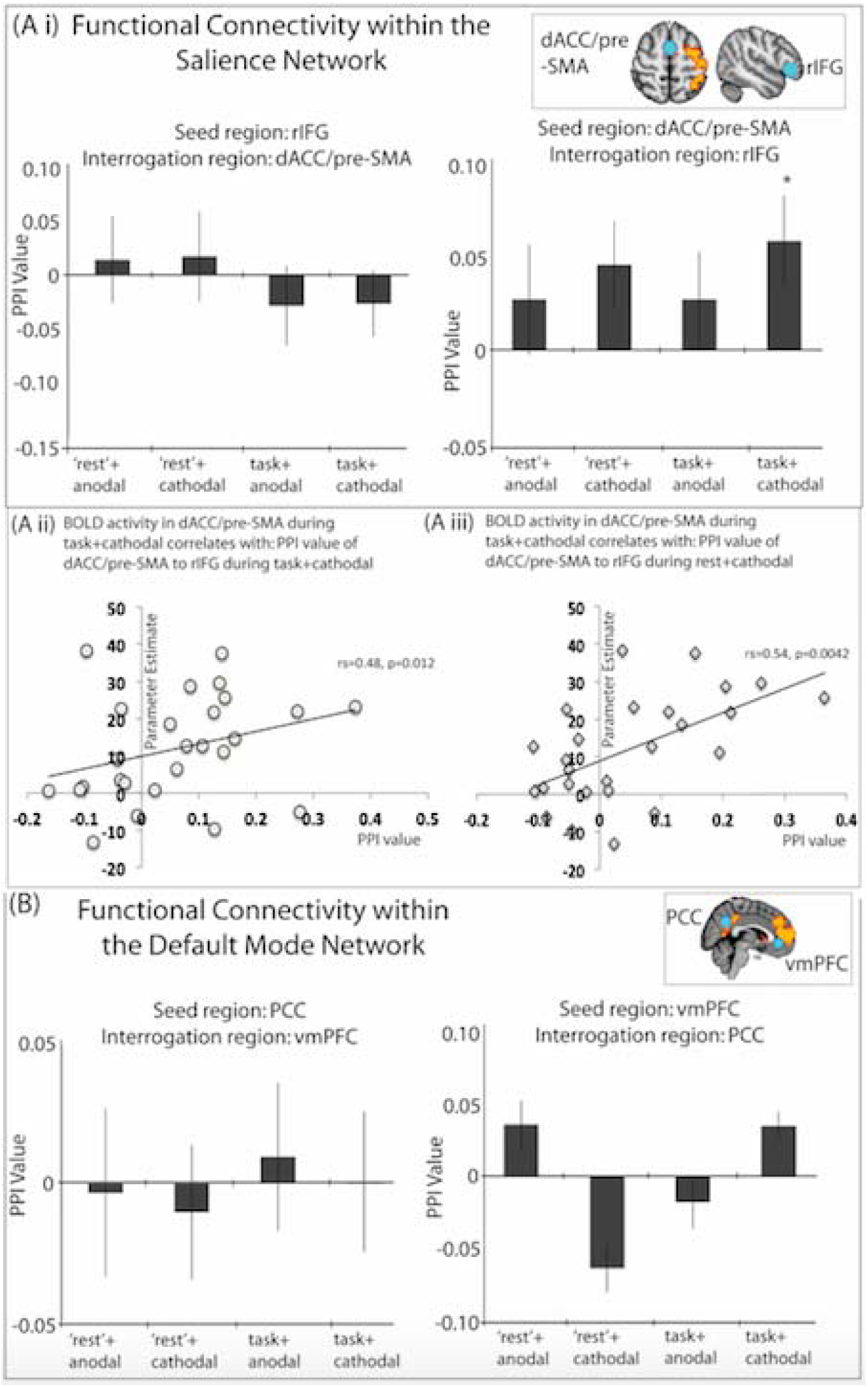
Functional Connectivity during different brain states and polarities. (Ai) Using dACC/preSMA as the seed, cathodal TDCS applied during task increases connectivity within the SN, though this does not survive Bonferroni correction for multiple corrections (* denotes p<0.05). Inset shows ROIs used for PPI analysis, superimposed on contrast Task: cathodal>anodal. (Aii) Correlation between SN PPI value during task+cathodal and (Aiii) correlation between SN PPI value during rest+cathodal with dACC/preSMA BOLD activity during task+cathodal. (B) There were no modulations of connectivity in the DMN under any condition. Inset shows ROIs used for PPI analysis, superimposed on contrast ‘Rest’: anodal>cathodal.

As stimulation of the rIFG led to large increases in BOLD signal within the dACC/pre-SMA during CRT performance we investigated whether the BOLD response in this region was related to the modulation of functional connectivity within the SN at ‘rest’ and/or during task. There was a significant correlation between the BOLD response in the dACC/pre-SMA during CRT+cathodal TDCS and the connectivity increase between the dACC/pre-SMA and rIFG, as measured with the PPI, during both CRT+cathodal TDCS (r_s_=0.48, p=0.012) (Figure 4Aii) and ‘rest’+cathodal TDCS (r_s_=0.54, p=0.0042) (Figure 4Aii). The latter relationship existed despite the functional connectivity between the dACC/pre-SMA and rIFG, as measured with PPI, not being significantly increased during ‘rest’+cathodal. There was no relationship between the PPI value during CRT+cathodal and the BOLD activity of dACC/pre-SMA during task without stimulation (r_s_=-0.08, p=0.69). Neither of these two relationships was seen for activity within the rIFG, where there was no significant modulation of regional brain activity by TDCS, nor was it observed for the relationship between PPI and dACC/pre-SMA during anodal stimulation, despite the significant modulation of dACC/pre-SMA activity observed.

Significant modulations of functional connectivity were not observed from the rIFG to the dACC/preSMA in either rest or CRT conditions, and no modulation of DMN connectivity was observed in any condition (Figure 4B).

### Behaviour

The effects of TDCS on regional brain activity and functional connectivity were seen in the absence of significant changes in behaviour and so interpretation of the neuroimaging results is not confounded by condition differences in CRT performance. A three level ANOVA with stimulation type as factors (sham/anodal/cathodal) did not show a main effect of stimulation on either CRT accuracy or any parameter of the exGaussian distribution for overall reaction times (all F<1, all p>0.05).

### Subjective experience

Prior to having combined TDCS and MRI, participants received 2 blocks of anodal and cathodal TDCS (15s each) in a randomised, blind order. Participants were asked to rate their sensation of ‘itching’, ‘pain’, ‘metallic taste’, ‘burning’, ‘anxiety’ and ‘tingling’ on a scale of 1-5 (1=nil, 2=mild, 3=moderate, 4=strong, 5=unbearable). There were no differences observed between average ratings given to anodal versus cathodal TDCS on all categories.

## Discussion

Until now, TDCS studies have treated underlying network state and polarity as independently acting factors. We conclusively show that widespread modulation of networks involved in cognitive control is achievable with TDCS, of even brief durations, when applied to the right inferior frontal gyrus (rIFG), both during task performance and at ‘rest’. This effect is dependent on the underlying state of the brain network and the polarity of stimulation. Furthermore, we demonstrate an interaction between TDCS polarity and cognitive network state, such that the same polarity TDCS caused distinct effects on the brain depending on whether subjects were engaged in a cognitively demanding task or were at ‘rest’. These findings are important both for interpreting previous studies, potentially explaining the variability of TDCS effects, and also for shaping the design of future stimulation studies.

Previous TDCS studies have mainly focused on the effects of stimulation on motor cortex, concluding that anodal TDCS increased neuronal excitability and cathodal TDCS reduced it (17) (18,19). However, it is unclear whether these results can be extrapolated to other parts of the cortex. Many cognitive studies have investigate the effects of TDCS on behaviour. However, relatively few (∼20%) have shown the expected distinction between anodal and cathodal stimulation (20). Only a small number of studies have used concurrent TDCS/fMRI to directly investigate physiological effects, and these have shown that anodal and cathodal TDCS are both capable of producing increases and decreases in cortical activity and connectivity (8,9,21). Our study extends this work by showing that the effects of TDCS polarity depends on the state of the network when it is stimulated, demonstrating that the effect of polarity is more nuanced than a simple dichotomy where anodal stimulation produces excitation of cortical activity and cathodal stimulation is inhibitory.

### Stimulation of the right inferior frontal gyrus modulates activity in large-scale cognitive networks

Brief stimulation of a single brain region, the rIFG, modulated activity in remote parts of two large-scale brain networks involved in cognitive control, the DMN and FPCN (13,22–24). The rIFG acts as a hub connecting many other cortical regions and is activated by a wide range of cognitive functions (25). It is thought to coordinate changes in activity across other cognitive control networks when switching between different task states (12,13,26,27). Hence, we reasoned that stimulating this region may produce widespread network changes in remote but connected brain regions, potentially accentuating the control mechanism exerted by the region.

We are unaware of any previous TDCS-fMRI study investigating the effects of rIFG stimulation. However, a small number of TDCS-fMRI studies applying TDCS to the primary motor cortex (M1) or dorsolateral prefrontal cortices (dlPFC) have shown effects distant from the site of stimulation (5–8,28,29). Additionally, a study comparing M1 and left dorsolateral prefrontal cortex TDCS, found that M1 TDCS modulated connectivity of sensorimotor networks, whilst TDCS to the DLPFC additionally modulated affective networks (30). However, the remote effects of TDCS and its interaction with task have not been systematically investigated for networks involved in cognitive control.

### Short durations of stimulation produced large physiological effects

Large changes in BOLD activity were seen after seconds, rather than minutes, of stimulation. This fits with in vitro animal studies that show applying cortical surface currents cause immediate changes in evoked potentials and spontaneous spike activity changes (17), and human studies showing that 4s of TDCS can produce changes in M1 excitability (Nitsche & Paulus 2000). Our study extend these findings by showing that rapid changes in activity of cognitive brain networks is possible with short durations of TDCS. TDCS has been shown to induce Ca_2+_ waves in astrocytes within seconds of application, suggesting that non-neuronal mechanisms might contribute to early neurobiological effects (32). However, purely non-neuronal mechanisms, such as direct effects of TDCS on brain haemodynamics, would be unlikely to have an interaction with task, which suggests that the effects of TDCS observed in our study reflect, at least in part, neuronal effects.

### The effects of stimulation depends on brain state

Our results clearly show that the physiological effects of TDCS are contingent on the current activity in that network. Distinct effects of the same type of TDCS were seen in a given network, as its activity varied with cognitive state. It has been shown that the spatial relationship between activated areas and deactivated areas is preserved across different brain states, and may reflect a homeostatic mechanism required for efficient brain function (33). TDCS appears to maintain this relationship, by enhancing both the deactivation, as well as the activation, associated with a given brain state, rather than causing global increases or decreases in activity.

Showing that the effects of TDCS are dependent on underying brain state, converges with animal work demonstrating that TDCS does not directly cause action potentials, but instead alters the probability of their occurrence (17)(34). Therefore, one would expect the effects of stimulation to vary depending on the populations of neurons active at that time.

A relationship between cognitive brain state and stimulation effects has been suggested by the small number of behavioural studies showing that manipulations of task difficulty can influence the behavioural modulations seen with TDCS (35–39). Transcranial alternating current stimulation (TACS) also shows effects on cortical network activity and connectivity that are dependent on the cognitive brain state (40–42). The link between the effects of stimulation and brain state has particularly important implications for clinical studies, since TDCS may produce distinct effects depending on whether it is applied during an active task or rest. Hence, attempts to use TDCS to enhance cognitive rehabilitation will need to carefully control the behaviour of a patient at the time of stimulation.

### The effects of stimulation polarity interact with underlying brain state

Our study demonstrates an interaction of underlying brain state and the polarity of stimulation on network activity. A similar interaction between task state and motor cortex excitability has been seen before, as assessed by motor evoked potential (MEP) size. MEP size was increased when anodal TDCS was given at rest, but was decreased if applied during task (43) However, to our knowledge, this interaction between brain state and TDCS polarity has not been seen before in cognitive networks or fMRI studies.

Current theories of how TDCS acts at the cellular level do not provide a simple explanation for this result. At the synaptic level, there is evidence that anodal and cathodal TDCS can have distinct effects on neurotransmitter levels. For example, the effects of anodal, but not cathodal TDCS, are abolished by NMDA_R_, voltage gated Ca_2+_ and Na_+_ receptor blockade (18). Additionally, a small number of studies have found that anodal TDCS decreases local GABA concentration and increases local Glutamine concentration, whereas cathodal TDCS decreases local Glutamine concentration (44–46). These changes could underlie the observed, and differential, effects of TDCS on local excitatory and inhibitory circuits (34,47–49). As local changes in the excitatory/inhibitory (E/I) balance are thought to produce changes in large-scale brain networks (50), this might provide a mechanism for the remote effects on network activity we observed. However, we show that both cathodal and anodal TDCS caused a change of BOLD response in the same direction relative to baseline, which cannot be explained by opposing effects on excitatory and inhibitory neurotransmitter levels.

Our results might be explained by a complex interaction between stimulation and cellular structure and orientation. In vitro and modelling studies demonstrate that the effect of TDCS on soma and dendrite polarisation is influenced by neuronal shape, cortical layer and the orientation of neuronal processes in the electrical field (51–54). A cortical region that is activated by a task will include subpopulations of neurons, some excitatory and some inhibitory, with different morphologies, orientations and occupying different cortical layers. As a result, different polarities of TDCS, which can really be considered as different directions of current flow, may activate different subpopulations of neurons within the same region. In addition, there is a complex relationship between alterations in excitatory/ inhibitory (E/I) balance and BOLD activity mediated by alterations in blood oxygenation and flow (55). For example, increased BOLD signal can increase secondary to activity of both excitatory and inhibitory circuits or increased activity of excitatory circuits with decreased activity of inhibitory circuits. Hence, an interaction of brain state and polarity may arise due to different subpopulations being activated under different combinations of task and polarity.

The differential effects of TDCS on neurons of different orientations also limits how much the effect of TDCS on cognitive networks can be predicted through extrapolating from results of motor cortex studies. Such studies use montages very different to ours, resulting in different patterns of current flow, and different electrical fields along the somatodendritic axis of neurons. Careful modelling studies, incorporating neuronal subpopulations, combined with in vivo electrophysiological measurements, would be informative in clarifying the interaction between polarity and neuronal orienation.

### Changes in network connectivity may explain changes in network activity

Cathodal TDCS applied during CRT performance increased functional connectivity (FC) within the SN. Cathodal TDCS also increased the BOLD response to the CRT task within the dACC/pre-SMA node of the SN. These two measures of network function correlated with each other, that is, SN connectivity correlated with SN activity during CRT with cathodal TDCS. The lack of a relationship between FC changes in the SN during CRT+cathodal and BOLD activity within the dACC/pre-SMA node during ‘rest’+cathodal, reduces the likelihood that the significant correlations observed are because of spurious, non-TDCS factors Of note is that SN connectivity at ‘rest’ with cathodal TDCS also correlated with the SN activity during CRT performance with cathodal TDCS.

Resting state studies show that neuronal activity and metabolism are correlated with FC as measured by fMRI (56–58), and that spatial patterns of resting state network BOLD activity and FC are also correlated (59). Previous studies have additionally found that resting state FC predicts task-induced network activity (60,61). This suggests that stimulation-induced changes in FC may underlie stimulation-induced changes in network activity, particularly in regions remote from site of stimulation.

### Limitations

We did not observe a behavioural effect of stimulation, which could be due to a number of reasons. The minimum duration of stimulation required to produce a behavioural effect is unclear, and it is possible that longer durations of stimulation might have produced an behavioural effect. Most cognitive studies studying behavioural modulation have used at least 10 minutes of stimulation. However, TDCS duration does not appear to have a linear relationship with either electrophysiological or behavioural measures (10,62). Dosing studies, particularly of non-M1 areas, will help to clarify the relationship between duration of stimulation and effects, particularly as duration may also interact with brain state and polarity.

We focused on rIFG stimulation and did not investigate the effects of stimulating other parts of the cognitive control system. Therefore, we cannot comment on whether the results we have seen are specific to the rIFG. Stimulation of other highly connected areas may also produce similar widespread network effects. Network hierarchy analyses, comparing multiple different TDCS targets, would be a sensible approach to test this hypothesis. Our experimental design also does not permit permit exploration of TDCS effects that may have persisted after the end of the stimulation. Studies of motor cortex suggest that the intra-stimulation and post-stimulation effects of TDCS on cortical excitability can be different (10,31). It is uncertain how long these post-stimulation effects last for and further work will be needed to clarify this issue for stimulation of cognitive control networks.

## Conclusion

The implications from our study are far-reaching. We demonstrate that widespread modulation of cognitive networks is readily achievable with TDCS, and that this effect is highly dependent on the underlying brain network state and polarity. The effect of stimulation, therefore, is an emergent property of the applied current in combination with the underlying brain state. Our results suggests many avenues for future investigations, strongly argues for the need for concurrent neurobiological assessment in cognitive TDCS studies, and has important implications for the translation of TDCS for clinical use.

## Methods

### Participants

We recruited healthy volunteers from the Imperial College Clinical Research Facility healthy volunteers list, with no history of neurological or psychiatric illness (n=26, 13F:13M) (mean age 38 years, s.d. 15.5 years). All volunteers gave written informed consent. The study conforms to the Declaration of Helsinki and ethical approval was granted through the local ethics board (NRES Committee London – West London & GTAC). All participants were naïve to TDCS.

### MRI acquisition and paradigm

A T1 and functional MRI (fMRI) sequences was acquired on a 3T Siemens Verio (Siemens, Erlangen, Germany), using a 32-channel head coil, with parameters similar to (42) (SI Methods).

FMRI was acquired whilst participants performed a blocked Choice Reaction Task (Figure 1 & SI Methods). Each run had 12 task blocks and 12 rest blocks, interspersed with brief periods of black screen. During each block, participants received anodal, cathodal or sham TDCS, resulting in a factorial design, consisting of 4 blocks of 6 possible conditions: rest+sham; rest+anodal; rest+cathodal; CRT+sham; CRT+anodal; CRT+cathodal. The order of the blocks was pseudorandomised but the same across all participants. Each participant performed 3 runs sequentially, with a brief 2-3mins rest between acquisitions to prevent fatigue.

Participants also performed a separate shorter blocked CRT, with no TDCS, prior to the TDCS-fMRI paradigm in order to determine the basic patterns of BOLD activity during task performance (previously described (63)).

### Delivery of transcranial direct current stimulation

Stimulation was delivered using a MR-compatible battery-driven stimulator (NeuroConn GmbH, Ilmenau, Germany) with a previously described circuit (42). The active electrode (round, 4.5cm diameter) was placed over F8 (10-20 EEG system), which corresponds to the pars triangularis, and the return over the right shoulder (rectangular, 5x7cm) (longitudinal axis parallel to the coronal plane, halfway between base of neck and acromion tip). A computation model confirmed that the peak electric field strength was over the rIFG (SI Methods).

### FMRI Analysis

Preprocessing was carried out as described. Briefly, fMRI images were brain extracted, motion corrected and registered to standard space in FMRIB’s Software Library (FSL (Smith, 2004; Jenkinson et al., 2012). FMRIB’s ICA-based Xnoiseifier (FIX (Griffanti et al., 2014; Salimi-Khorshidi et al., 2014)) was used to further remove noise components (SI Methods).

The fMRI-TDCS CRT was analysed with FSL’s FMRI Expert Analysis Tool (FEAT) as described (*SI Methods*). Mixed effects analyses of group effects were performed for each run separately using the FMRIB local analysis of Fixed Effects for the regressors of interest and the contrasts [task+anodal]>[task+sham], [task+cathodal]>[task+sham], [‘rest’+anodal]>[‘rest’+sham], [‘rest’+cathodal]>[‘rest’+sham]. The inverse contrasts were also run. A higher-level mixed effects (FLAME 1+2) analysis of group effects was performed to combine all runs and all participants. A third-level mixed effects (FLAME 1+2) analysis was performed to investigate: [‘rest’+anodal]> [‘rest’+cathodal] (and its inverse contrast) and [task+anodal]>[task+cathodal] (and its inverse contrast). The final Z statistical images were thresholded using a Gaussian random field-based cluster inference with a height determined by a threshold of Z>3.1 and a corrected cluster significance threshold of p=0.05.

### Functional Connectivity Analysis

Whole brain psychophysiological interaction (PPI) analyses (Friston et al., 1997)(15) were performed to assess the effect of TDCS and brain state on functional connectivity. We used the following regions of interest: rIFG and dACC/pre-SMA (forming the SN) and the PCC and vmPFC (forming the DMN). The other region within the network was used to extract the PPI values generated by the seed ROI. This allowed us to measure the PPI specifically between the nodes of each network (SI Methods).

### Statistical analysis of behavioural results

Statistical analyses of task performance were conducted using Matlab (Mathworks, Natick, MA) and R (www.r-project.org). We calculated accuracy (defined as the percentage of correct responses, and modelled individual overall reaction times (RT) and first RTs (the RT of the first trial within each block) with an exGaussian distribution (64).

## Acknowledgements

LML is supported by a Wellcome Trust Clinical Research Training Fellowship (103429/Z/13/Z). IV is supported by a Sir Henry Wellcome Trust Fellowship (103045/Z/13/Z) and receives funding from the NIHR Imperial BRC. AO is supported in parts by NIH grants MH110217 and MH111439. DJS is supported by a National Institute for Health Research (NIHR) Professorship (NIHR-RP-011-048). We would like to thank Dr Jonathan Howard for tireless and insightful technical and methodological assistance.

